# Core genome sequencing and genotyping of *Leptospira interrogans* in clinical samples by target capture sequencing

**DOI:** 10.1101/2022.04.29.490004

**Authors:** Linda Grillova, Thomas Cokelaer, Jean-François Mariet, Juliana Pipoli da Fonseca, Mathieu Picardeau

## Abstract

The life-threatening pathogen *Leptospira interrogans* is the most common agent of leptospirosis, an emerging zoonotic disease. However, little is known about the strains that are circulating worldwide due to the fastidious nature of the bacteria and its difficulty to be culture isolated. In addition, the paucity of bacteria in blood and other clinical samples has proven to be a considerable challenge for directly genotyping the agent of leptospirosis directly from patient material.

Here, to elucidate the genomic diversity of *Leptospira* circulating strains, hybridization capture followed by Illumina sequencing of the core genome was performed directly from 20 biological samples that were PCR positive for pathogenic *Leptospira*. A set of samples subjected to capture with RNA probes covering the *L. interrogans* core genome resulted in 72 to 13,000-fold increase in pathogen reads when compared to standard sequencing without capture. A SNP analysis of the genomes sequenced from the biological samples using 273 *Leptospira* reference genome was then performed in order to determine the genotype of the infecting strain. For samples with sufficent coverage (19/20 samples with coverage >8X), we could unambigously identify *L. interrogans* sv Icterohaemorrhagiae (14 samples), *L. kirschneri* sv Grippotyphosa (4 samples) and *L. interrogans* sv Pyrogenes (1 sample) as the infecting strain.

In conclusion, we obtained for most of our biological samples high quality genomic data at suitable coverage for confident core genome genotyping of the agent of leptospirosis. The ability to generate culture-free genomic data opens new opportunities to better understand the epidemiology and evolution of this fastidious pathogen.

## Introduction

Leptospirosis is a zoonosis of global distribution responsible for more than one million severe cases and 60,000 deaths with a higher incidence in tropical countries (Costa *et al*., 2015). The agent of leptospirosis belongs to the genus *Leptospira* which is composed of 68 species and more than 300 serovars (Vincent *et al*., 2019, Korba *et al*., 2021). The strains responsible for leptospirosis in humans or animals belong to one of the 8 pathogenic Leptospira species described until now. Among these pathogenic species, *L. interrogans* is the most frequently encountered worldwide (Guglielmini *et al*., 2019) and several studies have shown that strains belonging to the Icterohaemorrhagiae serogroup (*L. interrogans* serogroup Icterohaemorrhagiae), whose main reservoir is the rat, were responsible for the most severe forms of the disease (Hochedez *et al*., 2015, Tubiana *et al*., 2013, Herrmann-Storck *et al*., 2010, Christova *et al*., 2003).

Pathogenic leptospires are slow-growing bacteria that require a rich culture medium susceptible to contamination by other organisms. Isolation from biological samples is therefore tedious, especially since the bacteria can be present in low concentrations in blood and urine. During the course of the infection, the bacteria will be present in the blood during the first week after the onset of symptoms and concentration of *Leptospira* determined by qPCR ranged from 10^2^ to 10^6^ *Leptospira*/mL for the peak leptospiremia (Riediger *et al*., 2017, Agampodi *et al*., 2012). Leptospires are then found in blood in a decreasing number at 6-7 days after onset of symptoms until *Leptospira* nucleic acid is no longer detectable (Agampodi *et al*., 2012, Waggoner *et al*., 2015). *Leptospira* can also be detected in urine after symptom onset for a longer duration than blood. However, the bacteria may not be constantly present in the urine during the infection and both the concentration of bacteria and the duration of its excretion are poorly defined.

Identification of circulating strains in a particular region is essential to identify potential reservoir hosts and high-risk exposures and to establish appropriate control and prevention measures (development of vaccines, control of potential reservoirs, information for the general population, etc). Typing of the clinical isolates can also be important to identify strains or virulence factors associated with disease severity. However, as indicated above, culture isolation is challenging.

Given the value of whole genomes for phylogenetic, epidemiological and biological studies, there is an increased interest in obtaining genomes of pathogens from clinical samples. This is particularly true for pathogens that are found in low quantities in the host organism and are difficult to culture, as is the case for pathogenic *Leptospira*. Single alleles such as *rrs* (Cosson *et al*., 2014), *ligB* (Bourhy *et al*., 2010), *lflb1* (Perez & Goarant, 2010) and *secY* (Guernier *et al*., 2017, Mason *et al*., 2016, Grillová *et al*., 2020) can be directly amplified from the samples and sequenced for subtyping but this approach provides low-level resolution and does not allow discrimination among closely related species and strains. MLST schemes using several alleles can also be used for direct typing from clinical samples but this can result in incomplete allelic profiles (Mendoza & Rivera, 2021, Varni *et al*., 2018, Weiss *et al*., 2016) and it provides limited genetic information in the infecting strain. We recently developed a core genome MLST (cgMLST) scheme based on 545 genes that are highly conserved across the *Leptospira* genus (Guglielmini *et al*., 2019). This cgMLST allows the identification of pathogenic species, serogroups and closely related serovars. However, this highly discriminatory approach requires culture isolation of clinical strains. Direct sequencing from clinical samples results with high human host DNA contamination. Illumina sequencing of the cerebrospinal fluid of a patient with neuroleptospirosis showed, for example, that only 0.016% of the sequence reads correspond to the bacterial agent of leptospirosis (Wilson *et al*., 2014). Due to the low numbers of pathogens in clinical samples, several culture-independent genome sequencing methods have been recently developed using host depletion and/or microbial enrichment approaches. Targeted DNA enrichment which rely on the reference genome of the target bacteria has thus been used to retrieve DNA of bacterial pathogens such as *Chlamydia trachomatis* (Christiansen *et al*., 2014), *Mycobacterium tuberculosis* (Brown *et al*., 2015) and *Treponema pallidum* (Marks *et al*., 2018, Arora *et al*., 2016, Pinto *et al*., 2016) from clinical samples.

Here we describe a method utilizing biotinylated RNA probes designed specifically for *L. interrogans* DNA to capture the *Leptospira* core genomes defined by our cgMLST scheme (Guglielmini *et al*., 2019) directly from routine diagnostic samples (Figure 1). This study demonstrated for the first time the successful and accurate sequencing of *Leptospira* genomes directly from biological samples.

**Figure 1:**
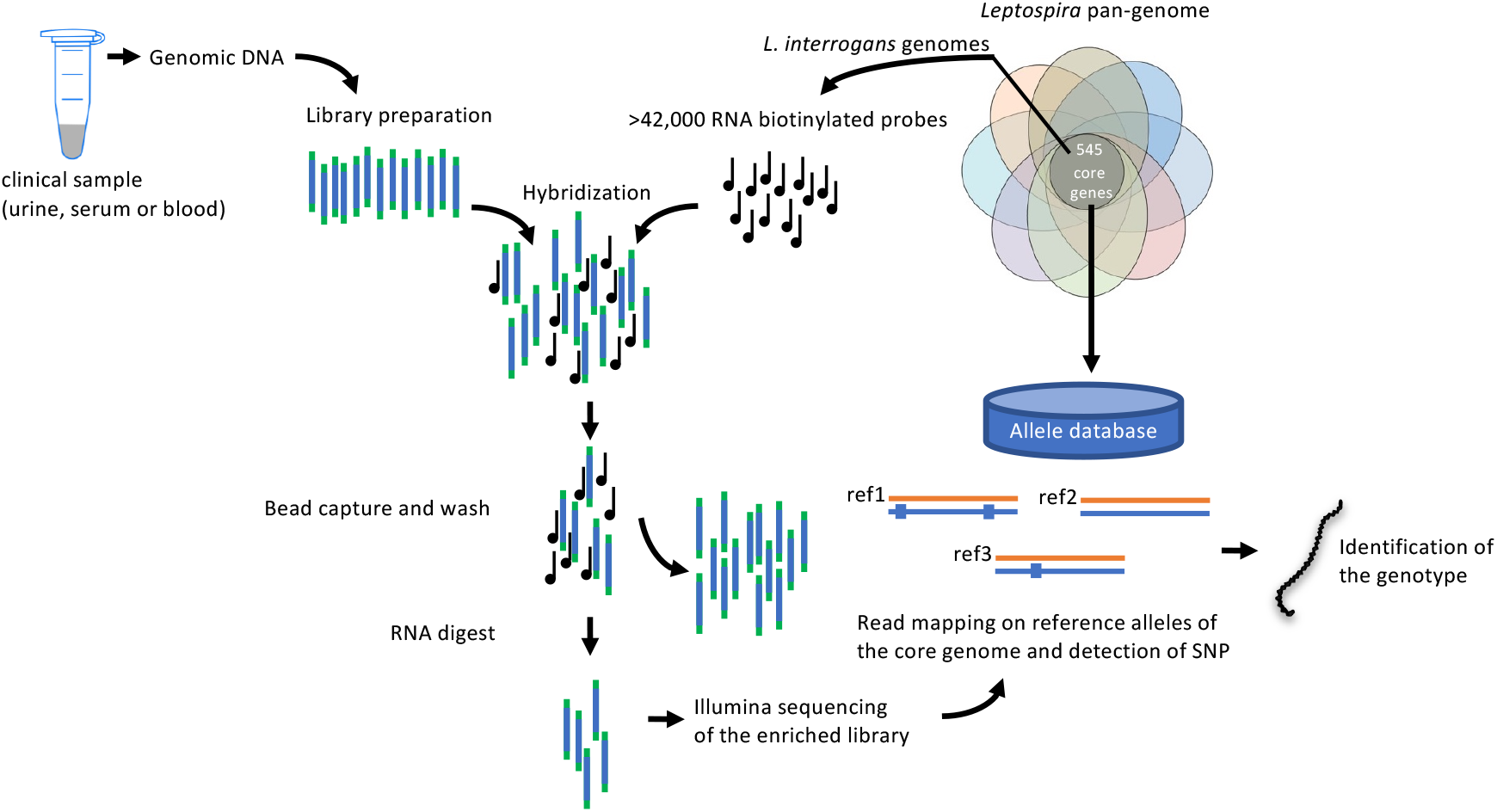
Schematic representation of the target enrichment method used in this study. For all samples, genomic libraries were prepared following the Sureselect Xt HS Target Enrichment System for Illumina. A library of pre-designed RNA probes based on 130 *L. interrogans* genomes was added to the genomic libraries. The target DNA (here the *Leptospira* core genome) hybridizes with the probes, but other DNA (human DNA) does not, allowing the uncaptured DNA to be removed during wash steps. The probe-bound DNA is then eluted and the enriched genomic libraries are collected for Illumina sequencing. Phylogeny and single nucleotide polymorphisms of the *Leptospira* sequences among patient samples is then performed by comparing with a database of 273 core genomes of pathogenic *Leptospira* strains to identify the infecting strain.

## Methods

### Samples

A total of 20 routine diagnostic samples (blood, serum or urine samples) tested positive by real-time PCR in the french National Reference Center (NRC) for Leptospirosis (Institut Pasteur) were analyzed in this study (Table 1). Total DNA was extracted using a DNeasy Blood and Tissue DNA extraction kit (Qiagen) Qiagen kit (Qiamp) and PCR was performed by real-time PCR using *lfb1* as a target (Bourhy *et al*., 2011). Sequencing of the PCR products of *lfb1* enable to identify the *Leptospira* species. *Leptospira interrogans* or the *L. interrogans* related species *L. kirschneri*-infected samples with a Ct value ≤38 were further selected for this study (Table 1). A written informed consent from patients was not required as the study was conducted as part of the routine diagnosis of the french NRC for Leptospirosis, and no additional clinical specimens were collected for the purpose of the study. Human samples were anonymized and human sequences were removed from the data before submission to the database. Collection of samples was conducted according to the Declaration of Helsinki.

**Table 1:** samples used in this study and identification of the infecting *Leptospira* strain.

| ID | source | Patient | Sex | Age | Ct value<sup>1</sup> | days<sup>2</sup> | nreads<sup>3</sup> | Bacteria (%)<sup>4</sup> | Species<sup>5</sup> | serogroup<sup>5</sup> | serovar<sup>5</sup> | strain<sup>5</sup> | Reference genome (id)<sup>6</sup> | number of SNP |
| --- | --- | --- | --- | --- | --- | --- | --- | --- | --- | --- | --- | --- | --- | --- |
| 1 | blood | A | M | 60 | 31 | unknown | 11,223,500 | 30 | *L. interrogans* | Icterohaemorrhagiae | Fiocruz | L1-130 | 246 | 3 |
| 2 | urine | B | M | 61 | 30 | 6 | 9,523,661 | 72 | *L. interrogans* | Icterohaemorrhagiae | Fiocruz | L1-130 | 246 | 6 |
| 3 | blood | C | M | 41 | 33 | unknown | 11,662,248 | 20 | *L. interrogans* | Icterohaemorrhagiae | Fiocruz | L1-130 | 246 | 3 |
| 4 | blood | D | M | 42 | 20 | 2 | 19,405,989 | 99 | *L. interrogans* | Icterohaemorrhagiae | Fiocruz | L1-130 | 246 | 3 |
| 5 | blood | E | M | 49 | 34 | 2 | 2,817,867 | 21 | *L. interrogans* | Pyrogenes | Pyrogenes | 2006006973 | 11 | 11 |
| 6 | serum | F | M | 40 | 34 | 4 | 6,936,356 | 13 | *L. kirschneri* | Grippotyphosa | Grippotyphosa | 201402975 | 117 | 6 |
| 7 | blood | G | M | 54 | 27 | 4 | 13,037,685 | 90 | *L. kirschneri* | Grippotyphosa | Grippotyphosa | 201402975 | 117 | 19 |
| 8 | blood | H | M | 20 | 32 | unknown | 8,739,223 | 65 | *L. interrogans* | Icterohaemorrhagiae | Fiocruz | L1-130 | 246 | 5 |
| 9 | blood | I | M | 69 | 29 | 3 | 12,665,561 | 65 | *L. interrogans* | Icterohaemorrhagiae | Fiocruz | L1-130 | 246 | 7 |
| 10 | blood | J | M | 74 | 28 | 0 | 12,989,647 | 94 | *L. interrogans* | Icterohaemorrhagiae | Fiocruz | L1-130 | 246 | 4 |
| 11 | blood | K | M | 36 | 32 | 4 | 9,589,816 | 90 | *L. interrogans* | Icterohaemorrhagiae | Fiocruz | L1-130 | 246 | 5 |
| 12 | urine | L | M | 24 | 31 | unknown | 10,827,346 | 65 | *L. interrogans* | Icterohaemorrhagiae | Fiocruz | L1-130 | 246 | 5 |
| 13 | urine | M | M | 73 | 31 | 5 | 11,947,042 | 90 | *L. interrogans* | Icterohaemorrhagiae | Fiocruz | L1-130 | 246 | 9 |
| 14 | urine | N | M | 39 | 35 | 8 | 29832,813 | 33 | *L. interrogans* | Icterohaemorrhagiae | Fiocruz | L1-130 | 246 | 5 |
| 15 | urine | O | M | 59 | 34 | unknown | 12,556,161 | 29 | *L. interrogans* | Icterohaemorrhagiae | Fiocruz | L1-130 | 246 | 5 |
| 16 | blood | P | M | 52 | 37 | 3 | 10,420,132 | 62 | *L. interrogans* | Icterohaemorrhagiae | Fiocruz | L1-130 | 246 | 6 |
| 17 | blood | Q | M | 44 | 25 | 5 | 1,350,830 | 98 | *L. interrogans* | Icterohaemorrhagiae | Fiocruz | L1-130 | 246 | 4 |
| 18 | blood | R | M | 22 | 38 | 4 | 1,0980,69 | 13 | *L. kirschneri* | ND | ND | ND | ND | ND |
| 19 | urine | S | M | 33 | 35 | 5 | 1,124,989 | 35 | *L. interrogans* | Icterohaemorrhagiae | Fiocruz | L1-130 | 246 | 2 |
| 20 | urine | T | M | 28 | 32 | 4 | 1,776,330 | 64 | *L. interrogans* | Icterohaemorrhagiae | Fiocruz | L1-130 | 246 | 4 |

<sup>1</sup> Ct value (Real-Time PCR Lfb1)
<sup>2</sup> days of sample collection since symptom onset
<sup>3</sup> number of Illumina reads
<sup>4</sup> proportion of reads in Bacteria (%), other reads correspond to human reads
<sup>5</sup> identification of the infecting strain based on the closest homologue of reference genomes
<sup>6</sup> identification number (id) of the reference genome in BIGSdb (www.bigsdb.pasteur.fr)

### SureSelect^XT^ target enrichment

A total of 42,117 custom-made SureSelect 120-mer RNA baits (total probe size 1,459 Mb) based on 130 *L. interrogans* genome sequences (Supplementary Table 1) spanning the 545 core genes previously defined for our cgMLST scheme (Guglielmini *et al*., 2019) were designed and synthesized by Agilent Technologies. For all samples, libraries were prepared following the Sureselect Xt HS Target Enrichment System for Illumina, from Agilent Technologies. For pre-capture library preparation, between 2 -200ng total gDNA were used. Briefly, samples were mechanically sheared using the Covaris E220 and the samples were repaired and the ends were a-tailed for barcoded Illumina adapters ligation. Ligated samples were amplified for 14 cycles and library quality was accessed using the Fragment Analyzer’s HS NGS migration kit. Libraries were captured individually, as recommended by Agilent. Captured libraries were pooled and sequenced using Illumina’s sequencing technology (Miniseq or Nextseq 500 sequencers). To access the efficacy of the capture method on patient’s samples, we also sequenced 5 libraries (samples 3, 4, 5, 7 and 8) before capture on an Illumina’s Miniseq sequencer.

### Sequence analysis

Prior to mapping, reads were trimmed from adapters. The variant calling pipeline described here after includes an additional trimming performed during the mapping step, which removes bases with Phred score below 30. An estimated depth of coverage was computing using the genome *L. interrogans* sv *Copenhageni* strain Fiocruz L1-130 as a reference. A taxonomic analysis based on Kraken2 (Wood *et al*., 2019) using human, bacterial and viral databases allowed us to classify 99% of the reads on average with a minimum of 97.2% in one sample.

We generated a database of 273 genome sequences of representative pathogenic *Leptospira* strains from our publicly accessible genome database online https://bigsdb.pasteur.fr/leptospira/, which is based on the software framework Bacterial Isolate Genome Sequence Database (Guglielmini *et al*., 2019). Our allele database includes genomes from 8 pathogenic species from 23 serogroups and 59 serovars isolated from patients from 40 countries (supplementary Table 2).

The variant calling analysis was performed with the variant calling pipeline (v0.10.0) from the Sequana project (Cokelaer *et al*., 2017) (https://github.com/sequana/variant_calling). The mapping step was performed with BWA (v0.7.17) (Li, 2013) and the variant calling was done with freebayes (v1.3.2) (Garrison & Marth, 2012). All VCF (Variant Calling Format) files (20 samples times 273 strains) were processed to gather the number of SNPs, INDELs, and MNPs in each sample and each strain. Scripts are available as notebooks on https://github/biomics-pasteur-fr/manuscript_capture_leptospira.

Variants were removed from subsequent analysis if one of the following conditions was met:

(i) a frequency of the alternate below 0.5 (minor variants), (ii) a strand balance below 0.2 (or >0.8) indicating an unbalance count of forward and reverse reads supporting the variant, (iii) a coverage below 10. In the case of sample 18 (highest Ct), we allowed the depth to be as low as 4.

### Nucleotide sequence accession number

The raw sequencing data have been deposited in array express with the accession number E-MTAB-11667.

## Results

Targeted enrichment has been applied to 20 biological samples, including blood, serum and urine samples from leptospirosis patients (Table 1). Library preparation, hybridization and subsequent enrichment were carried out on samples using the SureSelect Target Enrichment System (Agilent Technologies) (Gnirke *et al*., 2009) and custom designed RNA baits (Figure 1). To better evaluate the effectiveness of *Leptospira* capture, we compared the proportion of reads mapped to the *Leptospira* reference genomes with or without the SureSelect system for five samples (Figure 2). For samples prepared without the target-enrichment steps, the percentage of reads mapped to the *Leptospira* reference genomes are 0.0008% (sample 3), 1.36% (sample 4), 0.0008% (sample 5), 0.147% (sample 7), and 0.0127% (sample 8). For the same samples prepared using the SureSelect system, the percentage of reads mapped to the *Leptospira* genomes jump to 10%, 98%, 11%, 86% and 61%, respectively. Therefore, the capture increased the proportion of *Leptospira* by several order of magnitude (72 to 13,000).

**Figure 2:**
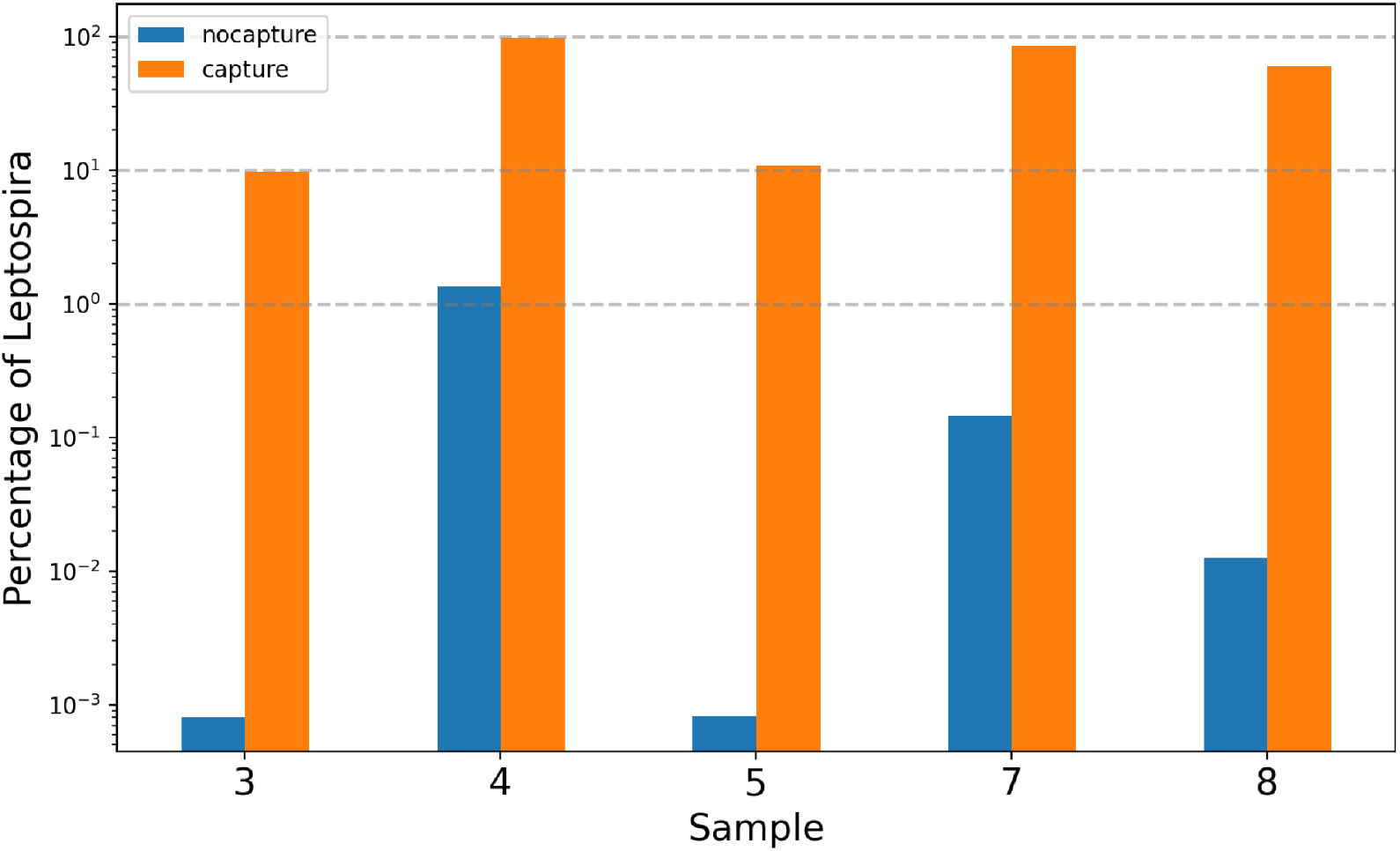
Percentage of *Leptospira* reads found in samples 3, 4, 5, 7, and 8 before (blue bars) and after hybridization capture using *Leptospira* RNA probes (orange bars).

The vast majority of bacterial reads correspond to the family *Leptospiraceae* (sup. Fig.1 and sup. Fig. 2); virus content is marginal (<0.1%) (sup. Fig. 2). In the 20 target enriched samples, the average depth of coverage was 590X with a range in 0.5 (sample 18) to 6000 X (sample 4) (sup. Fig. 3) hence leading to a large standard deviation of 1320. The coverage across the genomes was computed with the Sequana coverage tool (Desvillechabrol *et al*., 2018) to characterize more precisely the genomic variations in the different samples (sup. Fig. 4). Together with the average coverage shown in supplementary Figure 3, we can extract several key points. Most samples have coverage above 50X except for samples 3, 5, 6, and 18. The coverage of sample 3 and 5 remain large enough with 42X and 28X, respectively. Sample 6 has a low coverage of 8X. Finally, sample 18 is more problematic since its coverage is below 1X. Using *L. interrogans* sg Icterohaemorrhagiae (id246) as a reference (the closest strain for most samples), we can also look at the breadth of coverage (percentage of bases covered by at least one read) (sup. Fig. 4); it is above 99.5% in most samples except for sample 6, which has a breadth of coverage of 85%, and sample 18, which is about 3% only.

The Ct values of real-time PCR targeting the pathogen-specific target *lfb1* ranged between 20 and 38, which corresponds to 10e5 bacteria/µl to less than 1 bacteria/µl (Bourhy *et al*., 2011). A good correlation was observed between the Ct value and the proportion of *Leptospira* mapped reads and depth coverage (Figure 3). Thus, the 6 samples with more than 90% *Leptospira* reads (samples 4, 7, 10, 11, 13 and 17) have Ct values <32 (Table 1).

**Figure 3:**
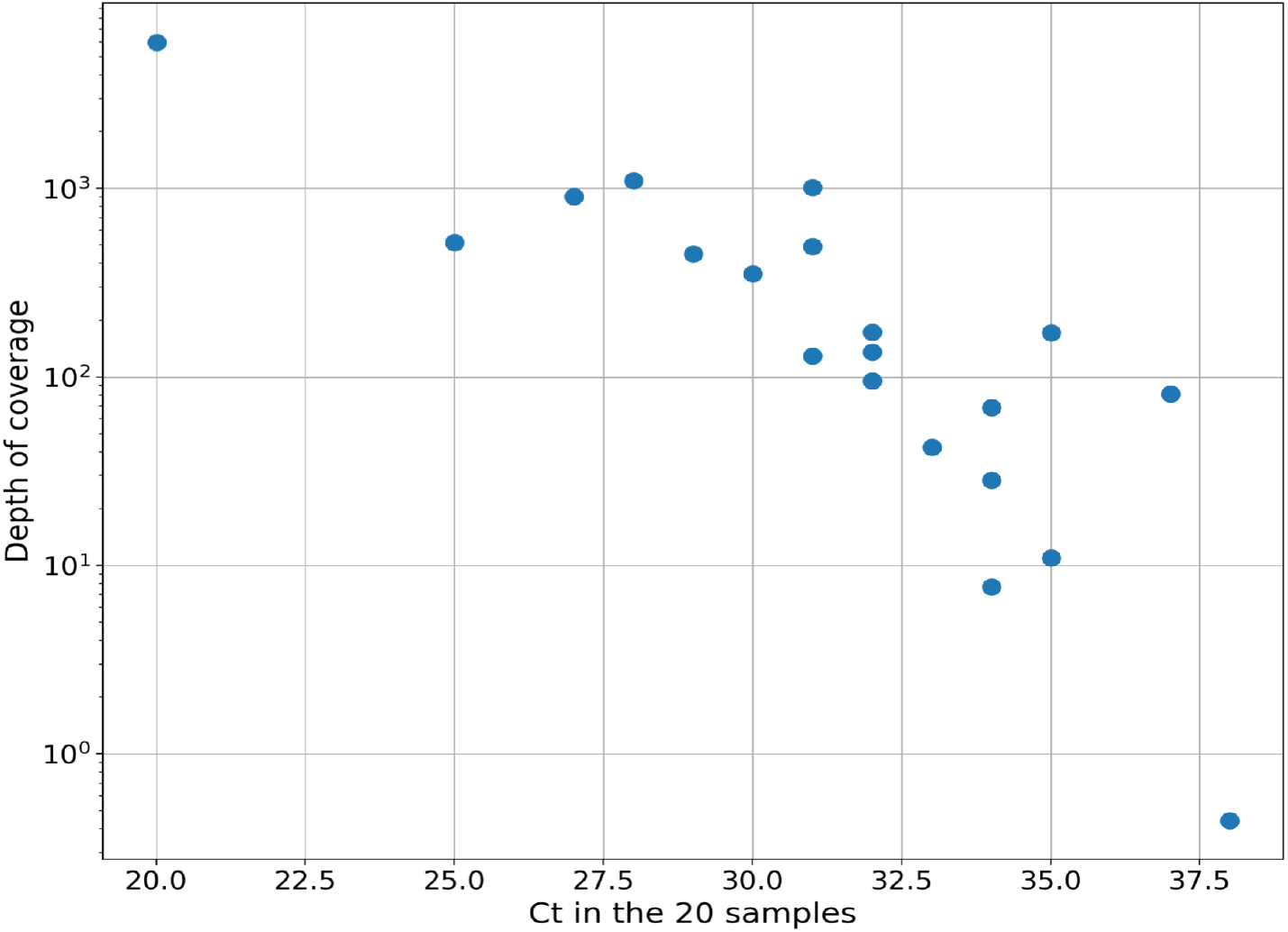
Final depth of coverage as a function of the measured Ct in the 20 samples. The depth of coverage is based on the *L. interrogans* core genome. The correlation coefficient between the depth of coverage and Ct values equals 0.76.

Variant calling approach was performed in order to identify each sample’s genotype. Given a database of 273 genomes of pathogenic *Leptospira* strains from different species, serogroups and serovars and originated from geographic areas (sup. Table 2), we searched for the closest genome to a given sample by minimizing the distance between the raw sequencing data of the sample and the *Leptospira* reference genomes. The distance used is the count of high-quality variants found in a given sample as compared to the different strains, as explained hereafter. Since the capture was designed with probes covering the core genomes only, we consider the core genome of the 273 strains; the average core genome length is 574.7kb +-8.5kb. Although coverage is uneven in some samples with the presence of spikes (excess of coverage in short regions; low frequency trend in sample 20) (sup. Fig.4), nevertheless it is generally high enough for a variant calling analysis. We first look at the number of SNPs. The distribution across all genomes and samples is highly variable with values ranging from 0 to 23,000 SNPs (average of 10,000). The SNPs count histogram across the 273 strains is shown in supplementary Figure 5 where 95% of the strains exhibit a count above 100 while a few strains have SNPs counts below 10 (Figure 4A and sup. Fig. 5).

**Figure 4:**
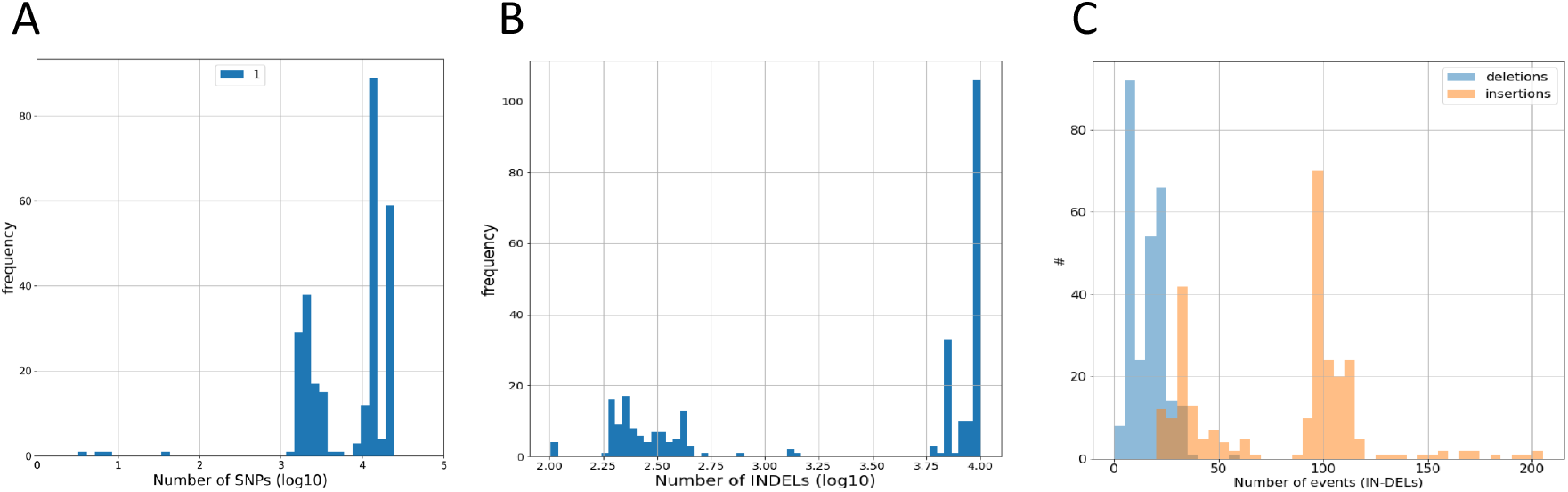
Analysis of multiple nucleotide polymorphisms. Histograms of Single-nucleotide polymorphisms (SNPs) (A), INDELs including multiple nucleotide polymorphisms (MNPs) (B), and insertions and deletions excluding MNPs (C) found in sample 1 across all 273 *Leptospira* genomes.

Interestingly, most of the examined samples have less than 10 SNPs when compared to the reference genomes of *L. interrogans* sv Icterohaemorrhagiae strain RGA (id97), *L. interrogans* sv Icterohaemorrhagiae strain Verdun (id106), and *L. interrogans* sv Copenhageni strain Fiocruz L1-130 (id246) (Table 2). These three strains are phylogenetically related (Figure 5) and they belong to *L. interrogans* sg Icterohaemorrhagiae. *L. interrogans* sv Copenhageni (id246) seems to be the strain that have the minimal number of SNPs in most samples (if we ignore samples 5, 6, 7 for which we are far from the correct strain, and sample 18 that has no SNPs due to low coverage). Sample 10 has 2 SNPs in id246 and only id106 has less with only 1 SNP (Table 2). Using the minimum number of SNPs as criteria for assignation, 14 out of the 19 samples were assigned to the genome *L. interrogans* sv Copenhageni (id246). Sample 10 was assigned to the genome *L. interrogans* sv Icterohaemorrhagiae (id106). Sample 5 was assigned to *L. interrogans* sv Pyrogenes (id11) and samples 6 and 7 to the genome *L. kirschneri* sv Grippotyphosa (id117) (Table 2). Sample 18 was excluded from the analysis due to a low average coverage of 1X. Indeed, with our stringent filter, less than 1 SNP was identified per strain. Nevertheless, we decreased the required depth to be as low as 4. In such case, the number of SNPs was about 20, on average, across all genomes (sup. Fig. 5). One had no SNPs (*L. mayottensis* sg Mini; id149) although 10-15 other strains had only 2-3 SNPs. These 10-15 strains are all close to each other in the phylogenetic tree (Figure 5) and belong to *L. kirschneri*. In particular, id117 has only 2 SNPs and was also the best hit in sample 6 and 7 (Table 2). In order to confirm those results, we also look at other type of variants: insertions, deletions and multiple nucleotide polymorphisms (MNP hereafter). The counts of insertions and deletions is summarized as number of INDELs hereafter. As shown in Figure 4BC, in sample 1, the number of INDELs varies from 100 up to 12,500. Using the minimum value across genomes, we found similar results as for the SNPs. For example, sample 1 has only 4 deletions and 98 insertions in id246. Sample 6 has several best hits. Indeed, id117 has 31 INDELs (29 insertions and 2 deletions) and only id110, id700 and id315 are equivalent. Interestingly, id117, id110 and id700 and id315 are next to each other in the phylogenetic branch. Using the minimum value across samples, we found similar results as for the SNPs. In sample 7, the id110, id117, id315 and id700 are also the best strains with 31, 32, 32 and 31 INDELs respectively. In sample 5, only 1 insertion was found in id11 confirming the SNPs result. All other samples have a minimum number of INDELs around 100. For example, sample 1 has 102 INDELs in id246, which is the same strain as in the case of the SNPs analysis confirming that sample 1 closest strain is id246. All other samples seem to have id246 for the closest strain as well (Table 2). Finally, we looked at MNPs. Sample 5 has 0 MNP only in id11. Samples 6 and 7 have a minimal number of 1 and 3 MNPs in id110, respectively. All other samples, which were close to id246, exhibit no MNPs in id246/id97/id106 (except sample 15 that has 1 MNP) (Table 2). Overall, the studies of SNPs/deletions/insertions/MNPs converge to a robust assignation of each sample.

**Table 2:** SNPs, INDELs and MNPs found in the 20 samples. The *Leptospira* sequences from samples was compared with a database of 273 core genomes of pathogenic *Leptospira* strains to find the closest homologue.

|  |  |  |  | samples |  |  |  |  |  |  |  |  |  |  |  |  |  |  |  |  |  |  |  |
| --- | --- | --- | --- | --- | --- | --- | --- | --- | --- | --- | --- | --- | --- | --- | --- | --- | --- | --- | --- | --- | --- | --- | --- |
| id | species | serovar | strain | 1 | 2 | 3 | 4 | 5 | 6 | 7 | 8 | 9 | 10 | 11 | 12 | 13 | 14 | 15 | 16 | 17 | 18 | 19 | 20 |
|  |  |  |  | Single Nucleotide Polymorphisms |  |  |  |  |  |  |  |  |  |  |  |  |  |  |  |  |  |  |  |
| 246 | interrogans | Copenhageni | Fiocruz L1-130 | 3 | 6 | 3 | 3 | 1462 | 4469 | 14924 | 5 | 7 | 4 | 5 | 5 | 9 | 5 | 5 | 6 | 5 | 1 | 2 | 4 |
| 106 | interrogans | Icterohaemorrhagiae | Verdun LP | 5 | 8 | 5 | 5 | 1765 | 4470 | 19852 | 7 | 9 | 6 | 7 | 7 | 11 | 7 | 7 | 8 | 7 | 1 | 1 | 6 |
| 97 | interrogans | Icterohaemorrhagiae | RGA | 6 | 9 | 6 | 6 | 1757 | 4436 | 19747 | 8 | 10 | 7 | 8 | 8 | 12 | 8 | 8 | 9 | 8 | 1 | 5 | 7 |
| 11 | interrogans | Pyrogenes | 2006006973 | 1588 | 1595 | 1555 | 1598 | 11 | 4449 | 19802 | 1555 | 1595 | 1596 | 1591 | 1592 | 1599 | 1590 | 1593 | 1586 | 1595 | 1 | 717 | 1588 |
| 117 | kirschneri | Grippotyphosa | 201402975 | 20825 | 20881 | 20417 | 21061 | 19884 | 6 | 19 | 20817 | 20916 | 20980 | 20833 | 20902 | 20968 | 20829 | 20730 | 20759 | 20917 | 0 | 9084 | 20792 |
|  |  |  |  | Deletions |  |  |  |  |  |  |  |  |  |  |  |  |  |  |  |  |  |  |  |
| 246 | interrogans | Copenhageni | Fiocruz L1-130 | 4 | 4 | 5 | 6 | 2 | 5 | 28 | 5 | 4 | 5 | 4 | 4 | 5 | 5 | 4 | 4 | 4 | 0 | 1 | 4 |
| 106 | interrogans | Icterohaemorrhagiae | Verdun LP | 4 | 4 | 5 | 6 | 2 | 5 | 27 | 5 | 4 | 5 | 4 | 4 | 5 | 5 | 4 | 4 | 4 | 0 | 1 | 4 |
| 97 | interrogans | Icterohaemorrhagiae | RGA | 5 | 5 | 6 | 7 | 3 | 5 | 28 | 6 | 5 | 6 | 5 | 5 | 6 | 6 | 5 | 5 | 5 | 0 | 1 | 5 |
| 11 | interrogans | Pyrogenes | 2006006973 | 7 | 7 | 8 | 10 | 0 | 5 | 29 | 8 | 7 | 9 | 7 | 7 | 8 | 8 | 7 | 7 | 8 | 0 | 1 | 7 |
| 117 | kirschneri | Grippotyphosa | 201402975 | 20 | 22 | 17 | 24 | 16 | 2 | 5 | 22 | 23 | 24 | 21 | 23 | 23 | 22 | 18 | 19 | 24 | 0 | 5 | 21 |
| 149 | mayottensis | ND | 200901116 | 10 | 15 | 7 | 25 | 6 | 4 | 32 | 11 | 12 | 20 | 10 | 13 | 16 | 11 | 10 | 11 | 11 | 0 | 0 | 10 |
|  |  |  |  | Insertions |  |  |  |  |  |  |  |  |  |  |  |  |  |  |  |  |  |  |  |
| 246 | interrogans | Copenhageni | Fiocruz L1-130 | 98 | 95 | 81 | 100 | 77 | 24 | 46 | 90 | 99 | 99 | 89 | 99 | 99 | 97 | 86 | 85 | 100 | 0 | 15 | 95 |
| 106 | interrogans | Icterohaemorrhagiae | Verdun LP | 99 | 96 | 82 | 101 | 78 | 24 | 94 | 91 | 100 | 100 | 90 | 100 | 100 | 98 | 87 | 86 | 101 | 0 | 15 | 96 |
| 97 | interrogans | Icterohaemorrhagiae | RGA | 97 | 94 | 80 | 99 | 76 | 23 | 92 | 89 | 98 | 98 | 88 | 98 | 98 | 96 | 85 | 84 | 99 | 0 | 14 | 94 |
| 11 | interrogans | Pyrogenes | 2006006973 | 101 | 97 | 83 | 102 | 1 | 26 | 94 | 90 | 101 | 101 | 90 | 100 | 100 | 99 | 89 | 87 | 101 |  | 14 | 97 |
| 117 | kirschneri | Grippotyphosa | 201402975 | 109 | 110 | 80 | 120 | 61 | 29 | 65 | 98 | 116 | 117 | 103 | 114 | 115 | 103 | 91 | 91 | 117 | 0 | 14 | 105 |
|  |  |  |  | Multiple nucleotide polymorphisms |  |  |  |  |  |  |  |  |  |  |  |  |  |  |  |  |  |  |  |
| 246 | interrogans | Copenhageni | Fiocruz L1-130 | 0 | 0 | 0 | 0 | 127 | 947 | 10912 | 0 | 0 | 0 | 0 | 0 | 0 | 0 | 1 | 0 | 0 | 0 | 0 | 0 |
| 106 | interrogans | Icterohaemorrhagiae | Verdun LP | 0 | 0 | 0 | 0 | 127 | 947 | 6364 | 0 | 0 | 0 | 0 | 0 | 0 | 0 | 1 | 0 |  | 0 | 0 | 0 |
| 97 | interrogans | Icterohaemorrhagiae | RGA | 0 | 0 | 0 | 0 | 126 | 938 | 6329 | 0 | 0 | 0 | 0 | 0 | 0 | 0 | 1 | 0 |  | 0 | 0 | 0 |
| 11 | interrogans | Pyrogenes | 2006006973 | 89 | 90 | 113 | 91 | 0 | 952 | 6403 | 89 | 90 | 91 | 90 | 92 | 93 | 90 | 97 | 91 | 89 | 0 | 42 | 89 |
| 117 | kirschneri | Grippotyphosa | 201402975 | 7032 | 7060 | 6987 | 7135 | 6702 | 1 | 3 | 7029 | 7064 | 7108 | 7038 | 7071 | 7086 | 7032 | 6990 | 7002 | 7042 | 0 | 2341 | 7044 |

**Figure 5:**
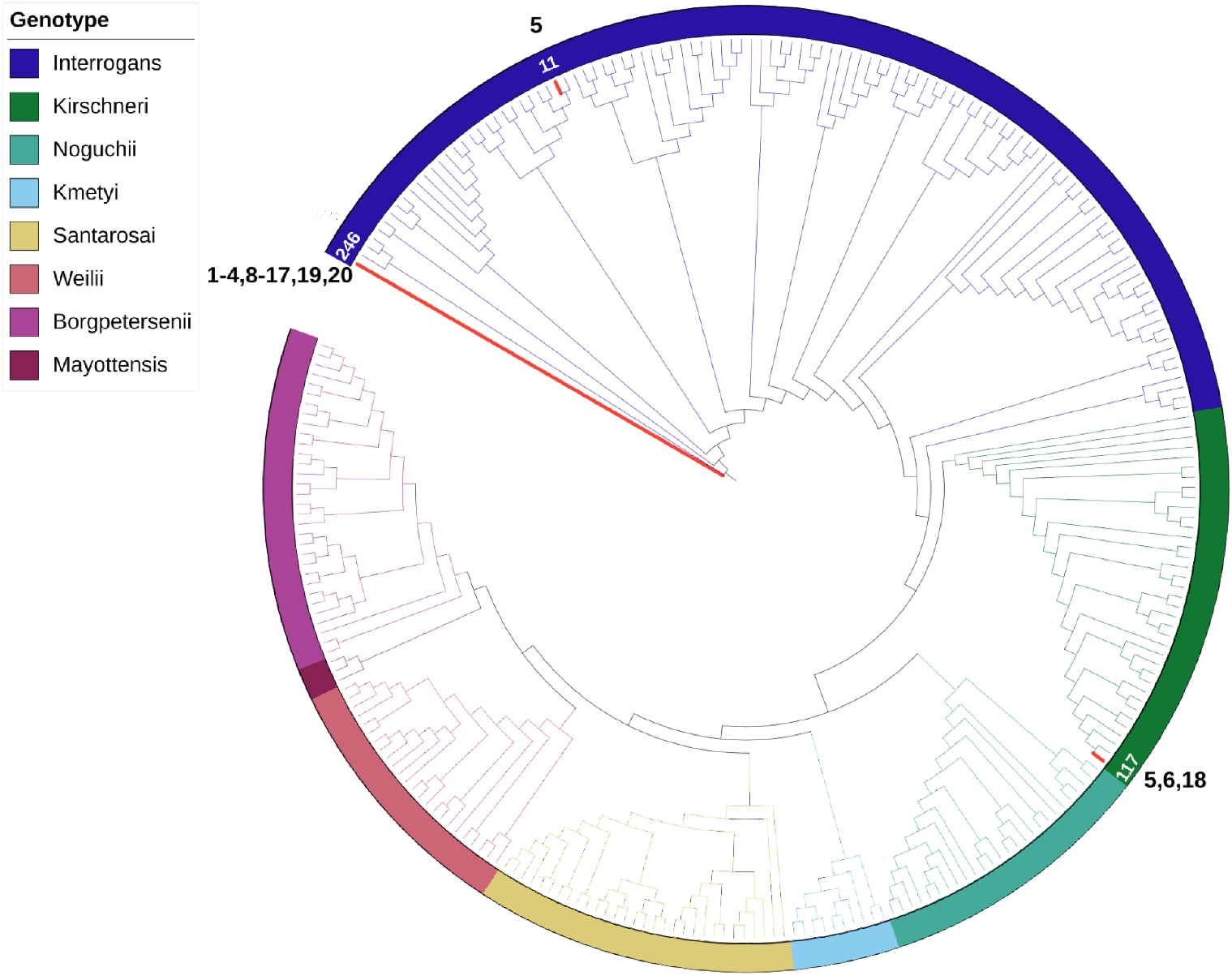
Phylogeny of *Leptospira* genomes and the sequenced samples. The *Leptospira* sequences from samples was compared with a database of 273 core genomes of pathogenic *Leptospira* strains (sup. Table 2) to find the closest homologue. Sample 1-4, 8-16, 17, 19, 20 have minimal number of single nucleotide polymorphisms (SNPs), deletions, insertions and multiple nucleotide polymorphisms (MNPs) on id246 (*L. interrogans* sv Icterohaemorrhagiae). Sample 6 and 7 and 18 are close to id117 (*L. kirschneri* sv Grippotyphosa) and Sample 5 is close to id11 (*L. interrogans* sv Pyrogenes). The phylogenetic tree was generated using multiple alignment (mafft v7.490) and RaxML software to infer the underlying phylogenetic tree. Visualisation was performed via the iTOL web service (https://itol.embl.de).

## Discussion

Genomics studies are proving key to the characterization of pathogen diversity and pathogenicity, yet the fastidious growth of *Leptospira* and the low abundance of *Leptospira* in clinical samples has presented a challenge for such studies. Thus, it can take up to four months of incubation for a primary culture to become positive (Wuthiekanun *et al*., 2007, Jayasundara *et al*., 2021). In addition, some *Leptospira* serovars require some additional culture media supplements for their growth (Hornsby *et al*., 2020). Our data demonstrate, for the first time, the suitability of target capture technology for purifying very low quantities of *Leptospira* DNA from complex DNA populations where the host genome is in vast excess. The successful enrichment of *Leptospira* DNA is shown by the significant increase in the ratio of bacteria:human DNA post-hybridization in a subset of samples. Ct was strongly correlated with capture efficiency. In six samples with a high leptospiral burden (qPCR Cts between 20 and 31 or 10e5 bacteria/µl to approximately 10e2 bacteria/µl), *Leptospira* reads accounted for > 90% of the total reads. Targeted enrichment has been applied to blood, serum and urine samples from leptospirosis patients a few days after symptom onset (0-8 days, average of 4 days) showing that the method can be applied to routine diagnostic samples. We show that enrichment of *L. interrogans* reads provides sequencing data that match the quality and quantity of data obtained via sequencing from culture with coverage above 50X for 16 out of the 20 samples, providing an opportunity to compare *Leptospira* strains from routine diagnostic samples with greater resolution than previously possible. Today this approach is still relatively expensive, currently costing approximately $300 per sample in our laboratory, but as next-generation sequencing costs continue to decline, this approach should be more affordable and accessible. To reduce the cost in future studies, samples can be barcoded and pooled before enrichment thus enabling multiplexing of hybridization reactions. This approach was already proven to decrease the price significantly (Marks *et al*., 2018, Beale *et al*., 2019).

The use of specific probes for *L. interrogans* is justified by the cosmopolitan nature of this species which is found worldwide (Guglielmini *et al*., 2019). The species *L. interrogans* also hosts the most pathogenic serovars such as the ones belonging to the Icterohaemorrhagiae serogroup (Hochedez *et al*., 2015, Tubiana *et al*., 2013, Herrmann-Storck *et al*., 2010, Christova *et al*., 2003). Finally, *L. interrogans* is particularly appropriate for the use of target enrichment, as *L. interrogans* has a relatively well-characterized clonal nature and *L. interrogans* strains from different origins showed high genetic relatedness (Guglielmini *et al*., 2019). Previous phylogenetic analyses indicated that *L. interrogans* serovar Copenhageni and Icterohaemorrhagiae strains from distinct geographical regions are highly conserved along time (Santos *et al*., 2018). The specificity of the target enrichment probe sets was confirmed by our ability to specifically target *L. interrogans* (17/20) but, interestingly, we were also able to target *L. kirschneri* (3/20) which is the closest species phylogenetically to *L. interrogans* (Table 1).

More importantly, this enrichment method effectively captures regions of diversity in the *Leptospira* core genome, which enables precise molecular typing of infecting strains. Comparison of these assembled sequences to the pathogenic *Leptospira* reference core genomes revealed only a limited number of SNPs. It is quite remarkable to see that the number of SNPs may be as low as a few SNPs (e.g. 3 SNPs in sample 1 on strain Fiocruz L1-130) even for sample with low sequencing yield. Analyzing the SNPs or INDELs or MNPs independently also give coherent results leading to robust assignation.

In the near future, a custom synthesized RNA probe sets could be designed to span the entire chromosome of *L. interrogans*. This will provide insight on bias that may be introduced by culture as previously shown for the spirochete *Treponema pallidum* (Pinto *et al*., 2017), but also increase our understanding in strain genetic diversity and its impact on disease outcome.

## Acknowledgements

We thank the staff of the National Reference Center for Leptospirosis (Pascale Bourhy, Céline Lorioux, and Farida Zinini) for support and processing some of the samples.

