## Supplementary tables and figures for "Core genome sequencing and genotyping of *Leptospira interrogans* in clinical samples by target capture sequencing"

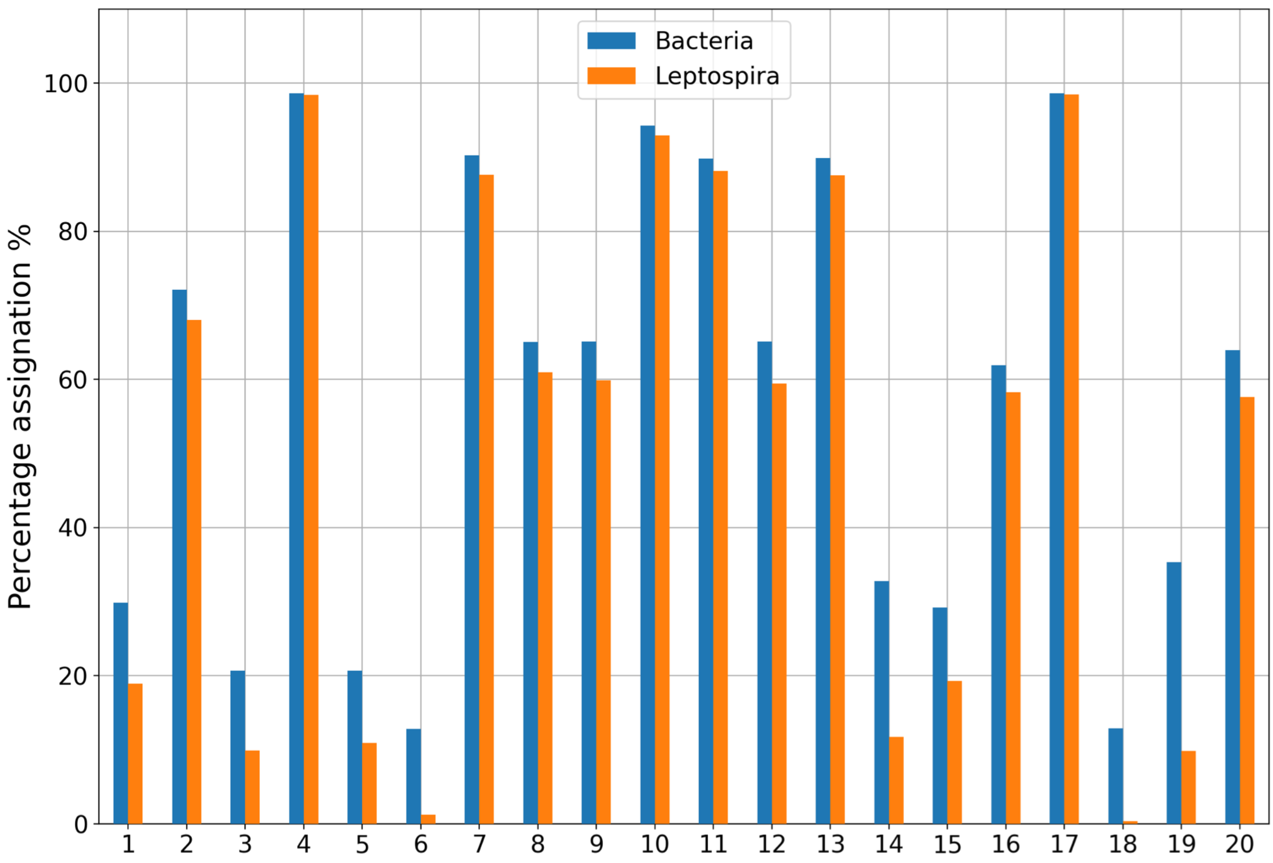


**Sup. Fig. 1** : Proportion of reads of the 20 samples assigned to bacteria genomes (blue bars) and proportion of reads assigned specifically to the *Leptospiraceae* family (orange bars).

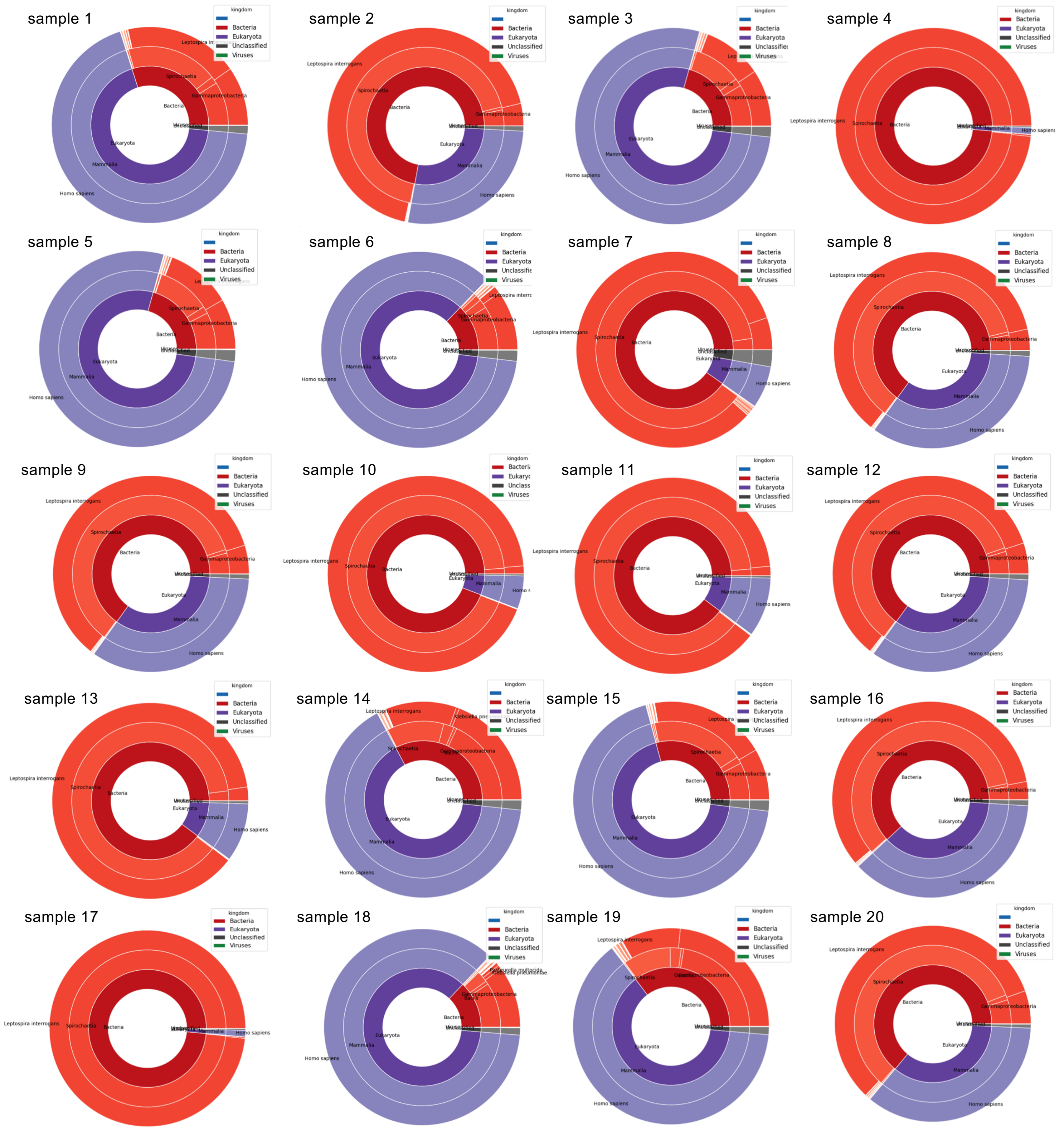


**Sup. Fig 2** : Taxonomic assignation of the reads of the 20 samples by Kraken2

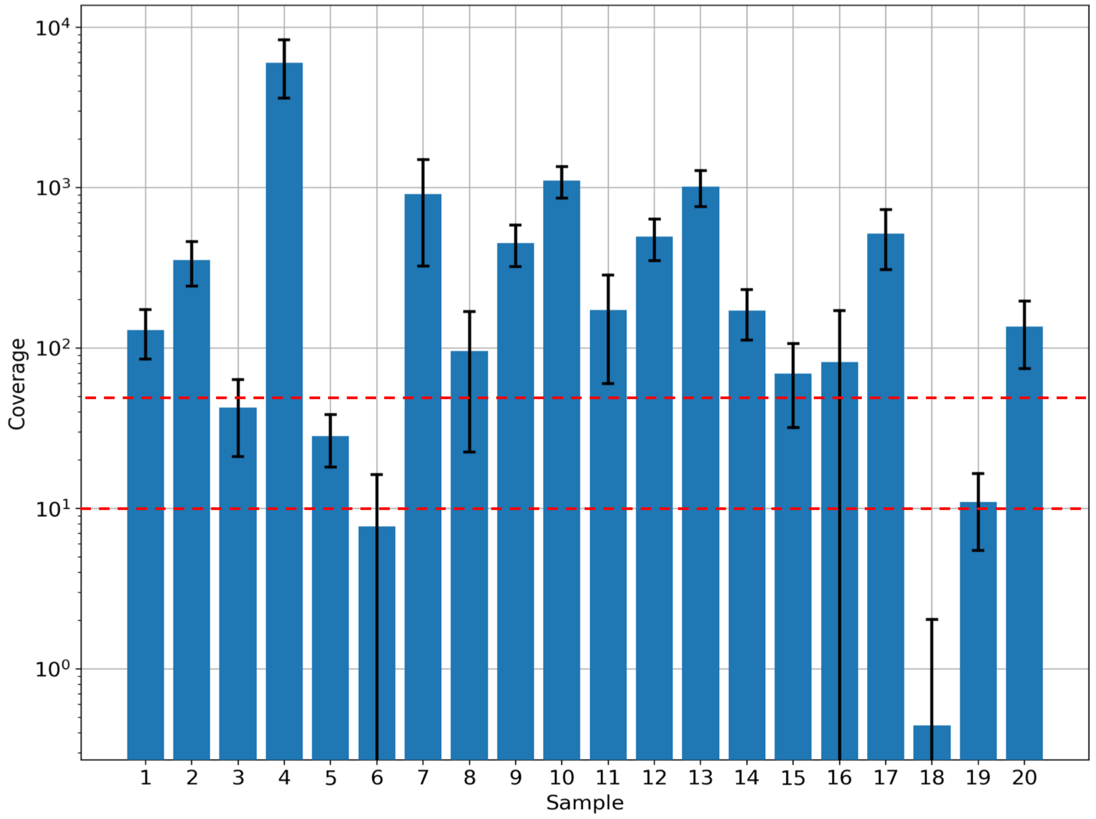


**Sup. Fig. 3 :** Coverage depth of the *L. interrogans core* genome across samples. The coverage depth on average for all samples is above 50X (top dashed red line) in most samples (sample 3 coverage equal 42X). Sample 5 and 6 have low coverage of 28X and 8X. Sample 18 coverage is below 1X. Black bars give the ±1 standard deviation.

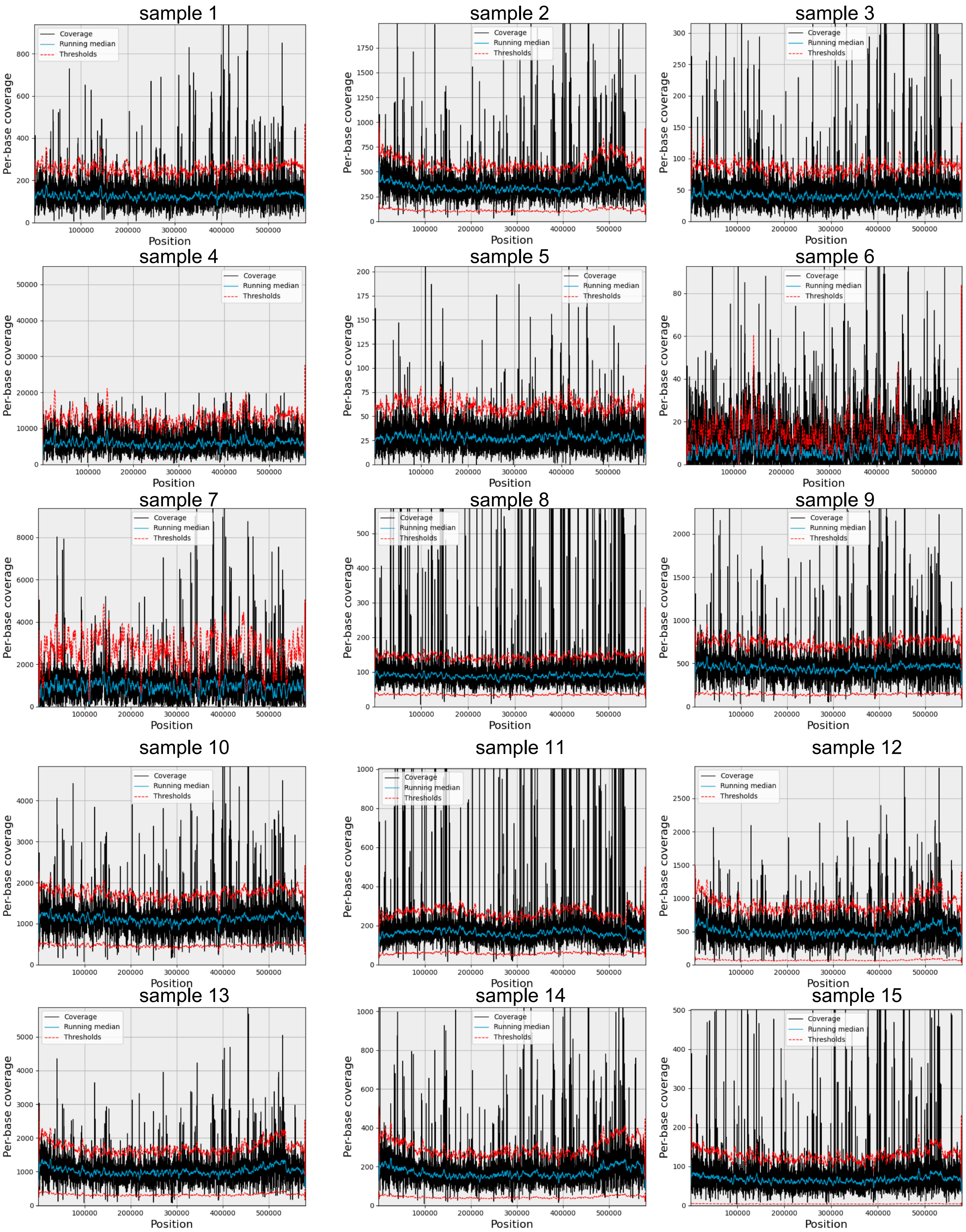
sample 1
sample 2
sample 3
sample 4
sample 5
sample 6
sample 7
sample 8
sample 9
sample 10
sample 11
sample 12
sample 13
sample 14
sample 15
Coverage
Running median
Thresholds
Per-base coverage
Position

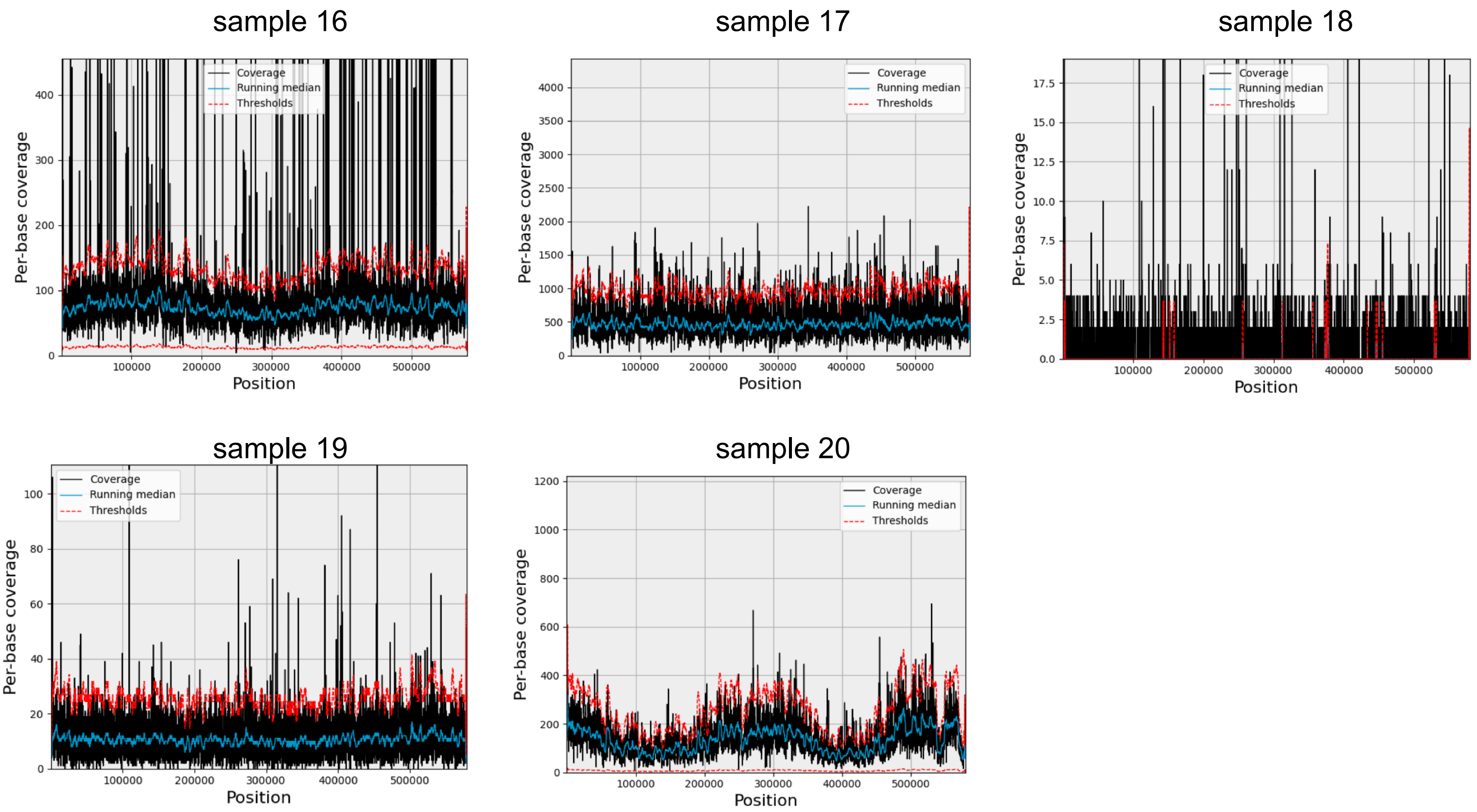


**Sup. Fig. 4** : Depth coverage of the 20 samples across the *L. interrogans* core genome

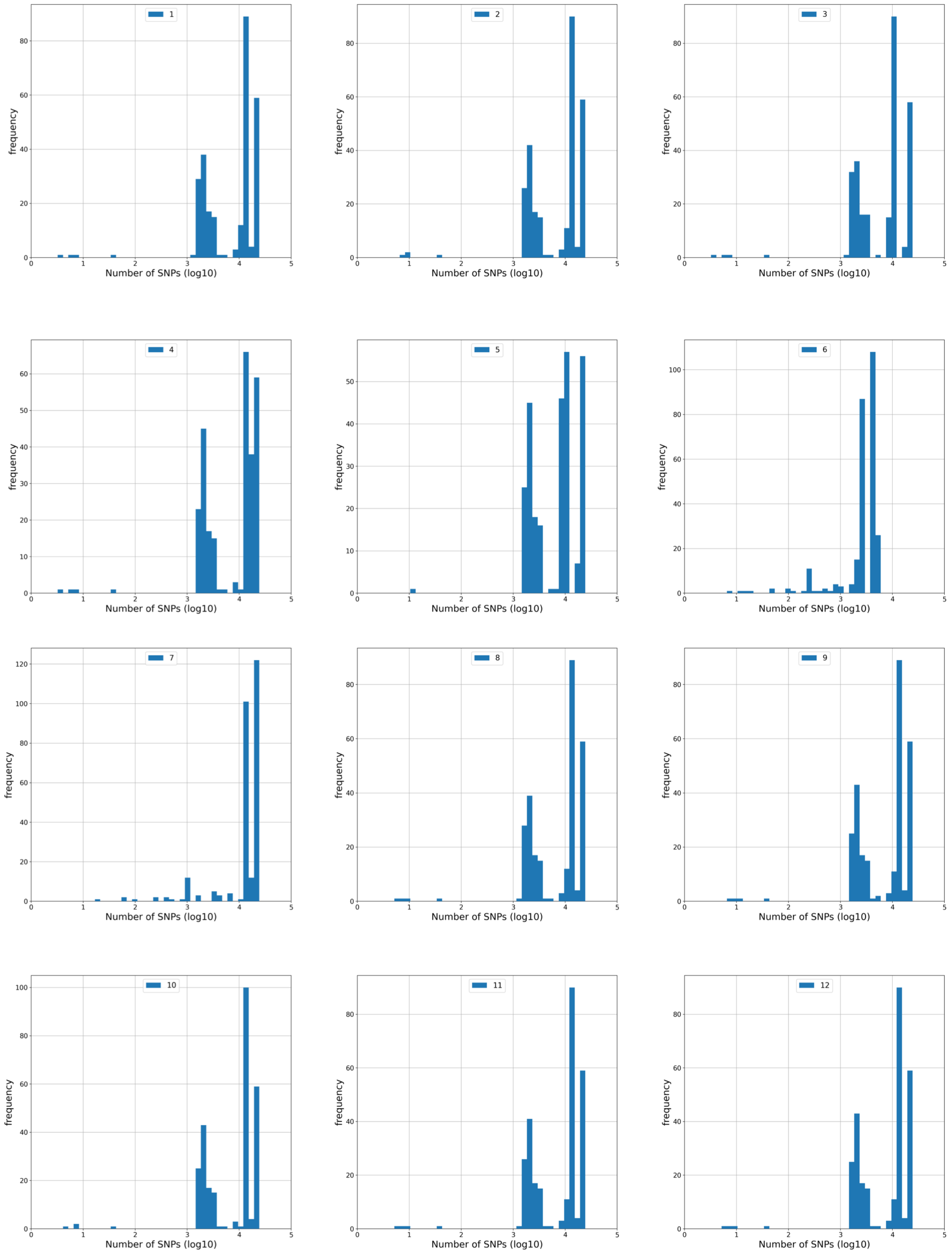

1
2
3
4
5
6
7
8
9
10
11
12
frequency
Number of SNPs (log10)

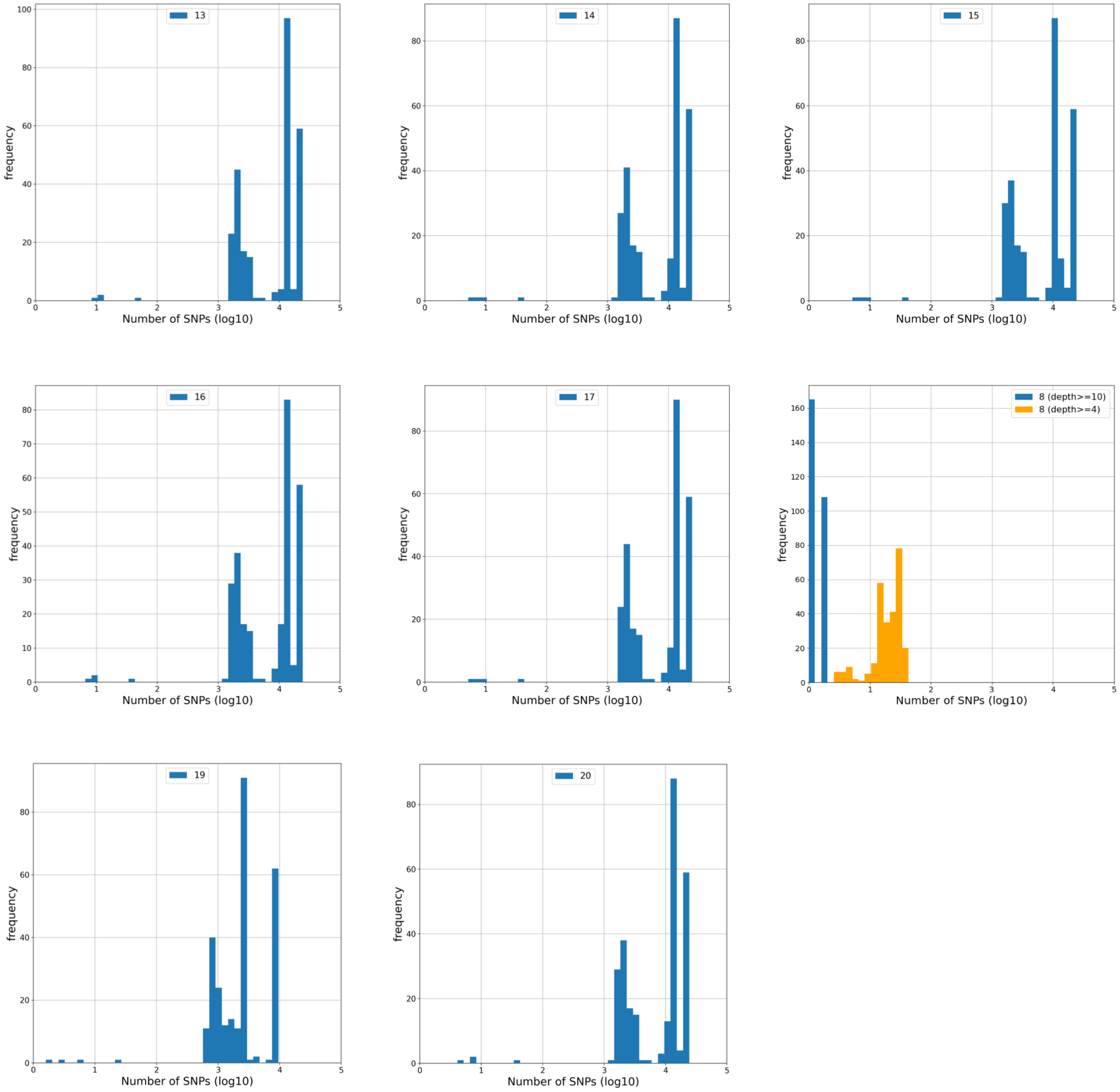


**Sup. Fig. 5** : Histogram of SNP counts found in the 20 samples across all 273 genomes.

For each of the 20 samples, we called variants using the 273 references of *Leptospira* (sup. Table 2) independently. For each genome, we obtained a set of SNPs that were filtered out to remove low quality variants (frequency below 10, uneven strand balance). Reference genomes distant from a sample led to thousands of SNPs while closely-related genomes led to less than 10 SNPs. Minimizing the count gives the closet genome to a given sample. Number of SNPs found in the 20 samples across all 273 genomes. Most genomes have more than 10,000 SNPs while only a few exhibit SNPs below 100.
